# Mirror bisulfite sequencing — a method for single-base resolution of hydroxymethylcytosine

**DOI:** 10.1101/019943

**Authors:** Darany Tan, Tzu Hung Chung, Xueguang Sun, Xi-Yu Jia

## Abstract

While the role of 5-methylcytosine has been well studied, the biological role of 5- hydroxymethylcytosine still remains unclear due to the limited methods available for single-base detection of 5-hydroxymethylcytosine (5hmC). Here, we present Mirror bisulfite sequencing detects 5-hydroxymethylcytosines at a single CpG site by synthesizing a DNA strand to mirror the parental strand. This semi-conservative duplex is sequentially treated with β- glucosyltransferase and M.SssI methylase. A glucosyl-5hmCpG in the parental strand inhibits methylation of the mirroring CpG site, and after bisulfite conversion, a thymine in the mirroring strand indicates a 5hmCpG site in the parental strand whereas a cytosine indicates a non-5hmC site. Using this method, the 5hmC levels of various human tissues and paired liver tissues were mapped genome-wide.

## INTRODUCTION

Since 5-hydroxymethylcytosines (5hmC) was discovered as an important epigenetic marker in 2009 (Kriaucionis and Heintz 2009; Tahiliani et al. 2009), a number of methods for genome-wide 5hmC profiling have been developed. Those methods can be classified into three main categories based on the approach used for 5hmC profiling: 1) enzymatic-based methods such as RRHP, HELP-GT assay and HMST-Seq (Petterson et al. 2014; Bhattacharyya et al. 2013; Gao et al. 2013), 2) affinity-based methods, like hMeDIP, hMeSeal and GLIB (Nestor and Meehan 2014; Song et al. 2011; Pastor et al. 2011), 3) bisulfite sequencing based methods, including TAB-Seq and oxBS-Seq (Yu et al. 2012; Booth et al. 2012). Although enzymatic-based methods allow straight-forward mapping of sequencing data, they are limited in their detection to the enzyme recognition site and may miss 5hmC sites which do not contain the corresponding enzyme recognition sequence. In contrast, affinity-based methods allow whole genome coverage, but they have relatively poor resolution and are biased toward 5hmC dense regions. In addition, neither of the two methods can quantitate 5hmC levels at single-base resolution. To address the issue, two bisulfite sequencing based methods, oxidative bisulfite sequencing (oxBS-seq) and TET-assisted bisulfite sequencing (TAB-Seq), had been developed. oxBS-seq utilizes potassium perruthenate to oxidize 5hmC into 5-formylcytosine (5fC), which is then susceptible to bisulfite conversion and sequenced as a thymine similarly to unmethylated cytosines. 5hmC sites are then quantified by subtracting oxBS-Seq data from traditional bisulfite sequencing (BS-seq). TAB-seq, on the other hand, first protects 5hmC through glucosylation before using a recombinant TET enzyme to oxidize all 5mC into 5-carboxylcyotsine (5caC). Therefore, after bisulfite conversion, only 5hmC sites would be sequenced as a cytosine. Although oxBS-seq and TAB-Seq can quantitate 5hmC at single-base resolution, both methods have limitations. oxBS-seq requires subtractive sequencing, which increases cost by doing two sets of bisulfite sequencing, while TAB-Seq relies on a highly efficient TET enzyme for complete oxidation of 5mC, which has been reported to be problematic at the end of genomic fragments (Hahn et al. 2015). Thus, a cost efficient and reliable method for 5hmC detection with single-base resolution is needed.

Here we report a novel method, Mirror Bisulfite Sequencing (Mirror-Seq), for the detection and quantification of 5hmC with base resolution precision at all CpG sites. This method utilizes highly efficient, commercially available enzymes, β-glucosyltransferase (βGT) and a CpG methylase, coupled with bisulfite conversion to interrogate 5hmC sites. Mirror-seq requires the synthesis of a single strand to mirror the parental strand, generating a semi-conservative duplex (Fig. 1a). This duplex is then sequentially treated with β-glucosyltransferase (βGT) and M.SssI CpG methylase. The presence of a glucosyl-5hmC on the parental strand inhibits methylation of the mirroring CpG site on the newly synthesized strand. After bisulfite conversion, the newly synthesized strands are selectively amplified and the cytosine mirroring the original 5hmCpG site is sequenced as a thymine. On the other hand, mirroring CpGs complementary to a non-5hmC site are effectively methylated and sequenced as a cytosine. By determining the methylation status of each CpG site in the newly synthesized strand, we can quantify the hydroxymethylation level on the original parent strand. Using this method, we were able to identify tissue-specific hydroxymethylated sites as well differentially hydroxymethylated sites in paired liver tumor samples.

**Figure 1.**
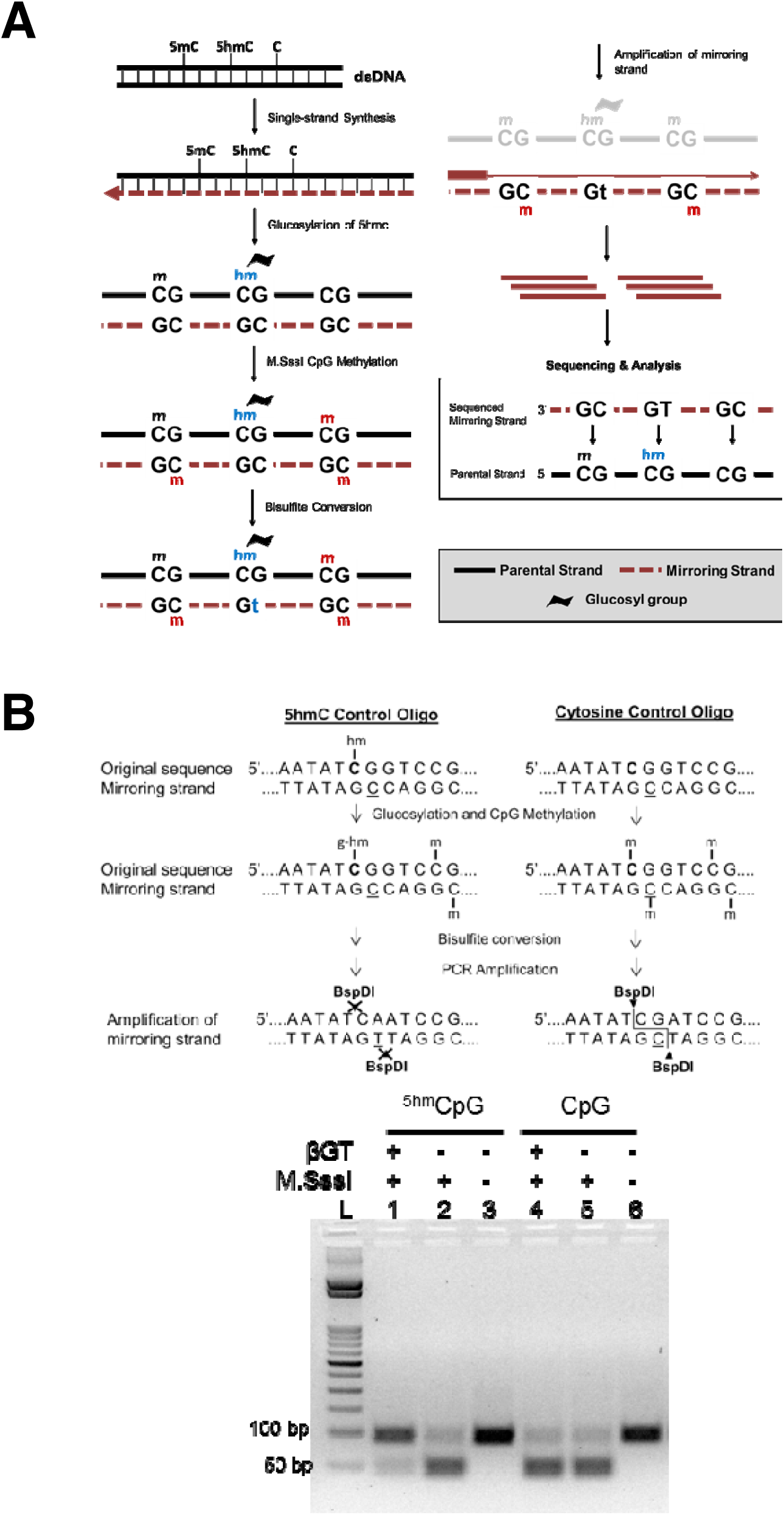
Using Mirror bisulfite sequencing to detect 5hmC in a CpG context. **(a)** Schematic overview of Mirror-seq workflow. A single-strand synthesis is used to generate a semi-conservative duplex in which the newly synthesized strand mirrors the parental strand. The duplex undergoes βGT and M.SssI methylation, consecutively. After bisulfite conversion, cytosines mirroring a 5hmCpG would be sequenced as a thymine while cytosines mirroring non-5hmCpG remain a cytosine. **(b)** Mirror-seq validation using a 100-mer oligo with a single 5hmCpG site (Lanes 1-3) or without any modified cytosines (lane 4-6). The amplicons were digested with BspDI to indicate the presence of a thymine or cytosine at the CpG site of interest (underlined). The presence of a thymine would result in an undigested amplicon (93 bp) where as a cytosine would result in two fragments (46 bp and 47 bp). L = 50 bp ladder (Zymo Research).

## RESULTS

To test the Mirror-seq methodology, we synthesized a 100-mer DNA oligo with a single 5hmCpG site at the 53^rd^ base (Fig. 1b). Using the 100-mer as a template, the reverse complementary strand was synthesized with unmodified dNTPs. The resulting dsDNA was sequentially treated with βGT and M.SssI methylase followed by conventional bisulfite conversion. After bisulfite treatment, the two strands were no longer complementary, and therefore, primers were designed to specifically amplify the newly synthesized strand. The resulting amplicon was then digested with BspDI, which recognizes the sequence ATˇCG^AT at the CG site of interest. Since the glucosyl-5hmC inhibited M.SssI methylation of the cytosine in the newly synthesized strand, the cytosine was converted into a thymine after bisulfite conversion, making it resistant to BspDI digestion (Fig. 1b, lane 1). The unglucosylated sample was efficiently methylated and retained the CG site, resulting in BspDI digestion (Fig. 1b, Lane 2). Untreated oligos had complete conversion of cytosines to thymines and, therefore, resistant to digestion (Fig. 1b, Lane 3). As a control, a 100-mer with the same sequence but with an unmodified cytosine instead of a hydroxymethyl group at the CG of interest was treated under the same conditions (Fig. 1b, Lanes 4-6). As expected, the amplicon was susceptible to BspDI digestion since there was no glucosyl-5hmC to inhibit methylation (Fig. 1b, Lanes 4). There appeared to be low amounts of undigested amplicon (93 bp), which may be due to incomplete BspDI digestion, incomplete CpG methylation, or impurity of the synthesized oligo. Nevertheless, these results demonstrated that a glucosyl-5hmC group inhibits M.SssI methylase activity, and by looking at the mirroring CpG site, we can determine the presence of a 5hmCpG.

The Mirror-Seq method can be adapted to a reduced representation bisulfite sequencing (RRBS) scale to allow deep, selective sequencing of the genome that is highly enriched for CpG islands (CGI). We generated Mirror-RRBS libraries using a pair of modified Illumina TruSeq adapters, and the single-strand extension was initiated by using a methylated primer specific to one of the ligated adapter (Supplementary Fig. 1). The methylated primer allowed the newly synthesized strand to be distinguished from the parental strand after bisulfite conversion and, therefore, to be selectively amplified and sequenced. After alignment and hydroxymethylation calling, the results were mapped back to the original strand since the sequenced strands were mirroring the parental CpG sites. Successful detection of 5hmC using Mirror-Seq is controlled by three key factors: (1) efficient glucosylation of 5hmC by βGT, (2) efficient M.SssI methylation of non-hydroxymethylated sites, and (3) efficient bisulfite conversion. We monitored those factors by spiking the genomic DNA prior to library preparation with four different PCR amplicons that represented different levels of 5hmC at a single CpG site: 100%, 50%, 5% and 0% (Supplementary table 1). The 5hmCpG site of each amplicon was used to evaluate the efficiency of Mirror-Seq in detecting various 5hmC levels. All of the other CpG sites were used to assess methylation efficiency while non-CpG cytosines were used to calculate the bisulfite conversion rate. Averaging the calculated rates across all four amplicons from different libraries, we obtained an average of 97.4 ± 0.3% methylation efficiency and 99.57 ± 0.03% bisulfite conversion rate (Supplementary table 2). The actual 5hmC levels calculated from the sequencing results were close to their theoretical value (Supplementary fig. 2) with an average difference of 1% between the theoretical and experimental value (Pearson’s R^2^ = 0.9988). These results confirmed the ability of Mirror-Seq for robust detection of 5hmC in the context of genomic DNA.

**Figure 2.**
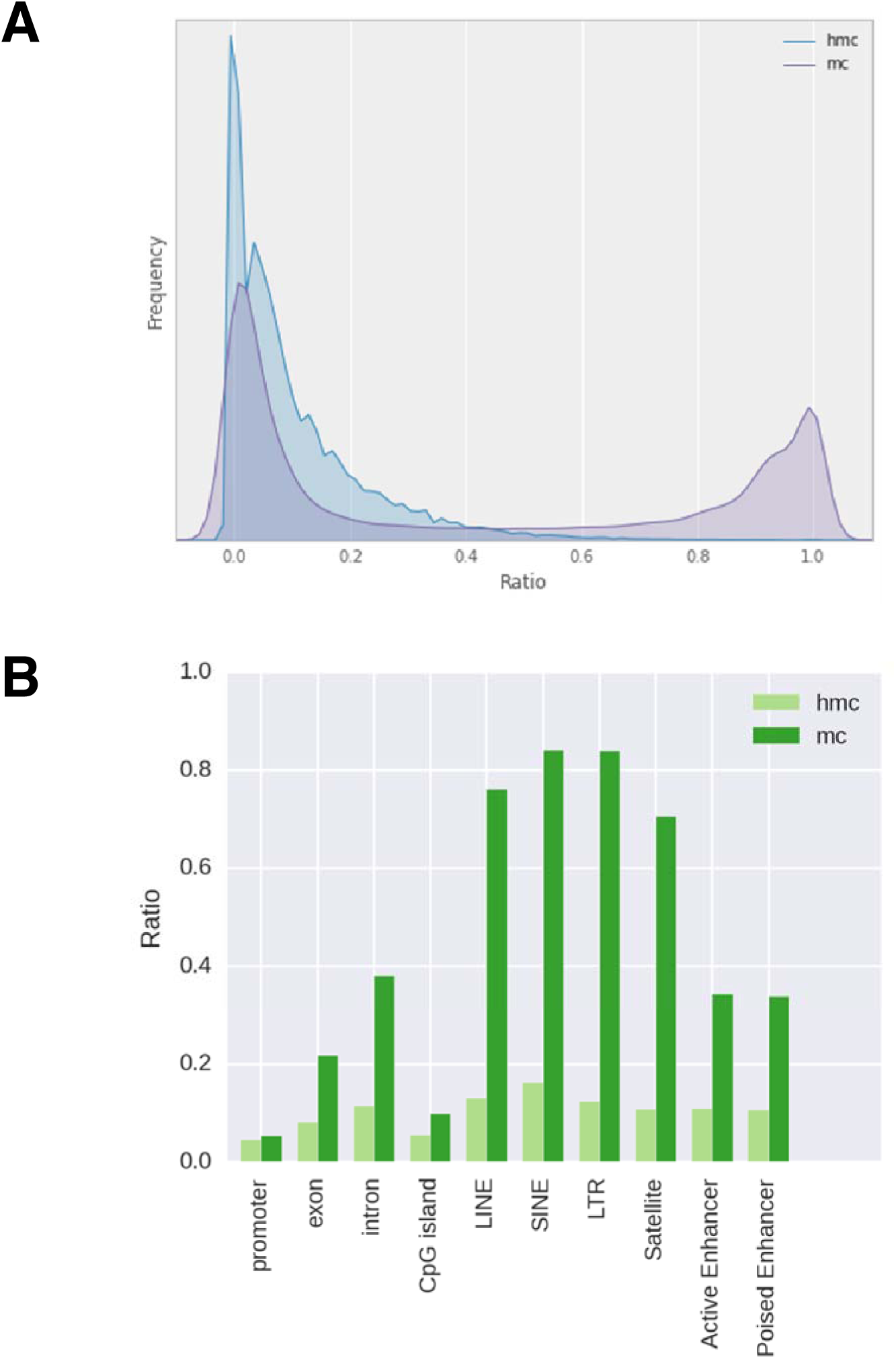
Scatter plots comparing technical replicates of Mirror-RRBS libraries. **(a, b)** The correlation between technical replicates for individual CpG sites were compared using 10 reads **(a)** and 50 reads cutoffs **(b)**. **(c,d)** Regional correlation analyses were also performed for promoters **(c)** and CpG islands **(d)**. Each annotated region contained at least three CpGs (≥10X coverage), and the plotted values were averages of the 5hmC ratios for the region.

This library preparation method was then used to quantitatively map 5hmC in the human brain, which has been shown to have the highest hydroxymethylated DNA among different tissues (Li and Jiu 2011). To test the reproducibility, two replicates of Mirror-RRBS libraries were generated from 400 ng of human brain DNA (denoted as 400A and 400B) and sequenced using 50 base pairs paired-end reads with about 50 million reads each (Supplementary table 3). Approximately 4 million CpG sites were detected with an average of 17-19x read coverage. A RRBS library was also prepared and sequenced from the same sample to evaluate the abundance of 5mC; consistent with previous studies (Wen et al. 2014), we found that the majority of CpGs had low 5hmC abundance in comparison to 5mC (Fig. 2a). The genomic distribution of the two modifications was quite different as well; 5hmC sites were more enriched in the gene body and enhancer regions whereas 5mC sites were more enriched in DNA repeat elements and intergenic regions (Fig. 2b). Pairwise comparison between the two technical replicates, 400A and 400B, showed the Pearson’s coefficient was closely related to the sequencing depth when single CpGs were compared. The Pearson’s coefficient was only 0.57 using 10x reads cutoff but increased to 0.96 when using 150x reads cutoff (Fig. 3a,b). However, the number of CpG sites decreased from about 3 million to 7 thousand, indicating that the number of sites with higher read counts was limited. To test the possibilities of utilizing CpG sites with low read coverage, we performed regional correlation analysis by averaging all the 5hmC sites within the same annotated region, such as CpG island and gene promoter. Using 10X read cutoffs, we found that region analysis greatly increased the Pearson’s coefficient compared to single site analysis (Fig. 3c,d and Supplementary fig. 3). This indicated that 5hmC analysis can be performed using lower sequencing depth by doing region analysis if single site assessment is not required. In addition, three Mirror-RRBS libraries were prepared from different inputs of genomic DNA to examine the minimal input requirement. We found that the DNA input can be decreased to 100 ng without changes to the library preparation protocol while 50 ng required additional PCR cycles during the final amplification step to reach the same yield (Supplementary fig. 4). Libraries generated from 200 ng and 100 ng inputs were sequenced and compared to library 400A. The correlations between libraries generated from different inputs were similar in terms of the Pearson’s coefficient at each read cutoff (Supplementary fig. 5). This indicated that lower input is feasible for Mirror-seq library preparation without compromising the library quality, which is of great value for investigating precious samples such as stem cells, certain clinical samples, or selectively isolated cellular subpopulations (i.e. diverse neuronal cells from a whole brain sample).

**Figure 3.**
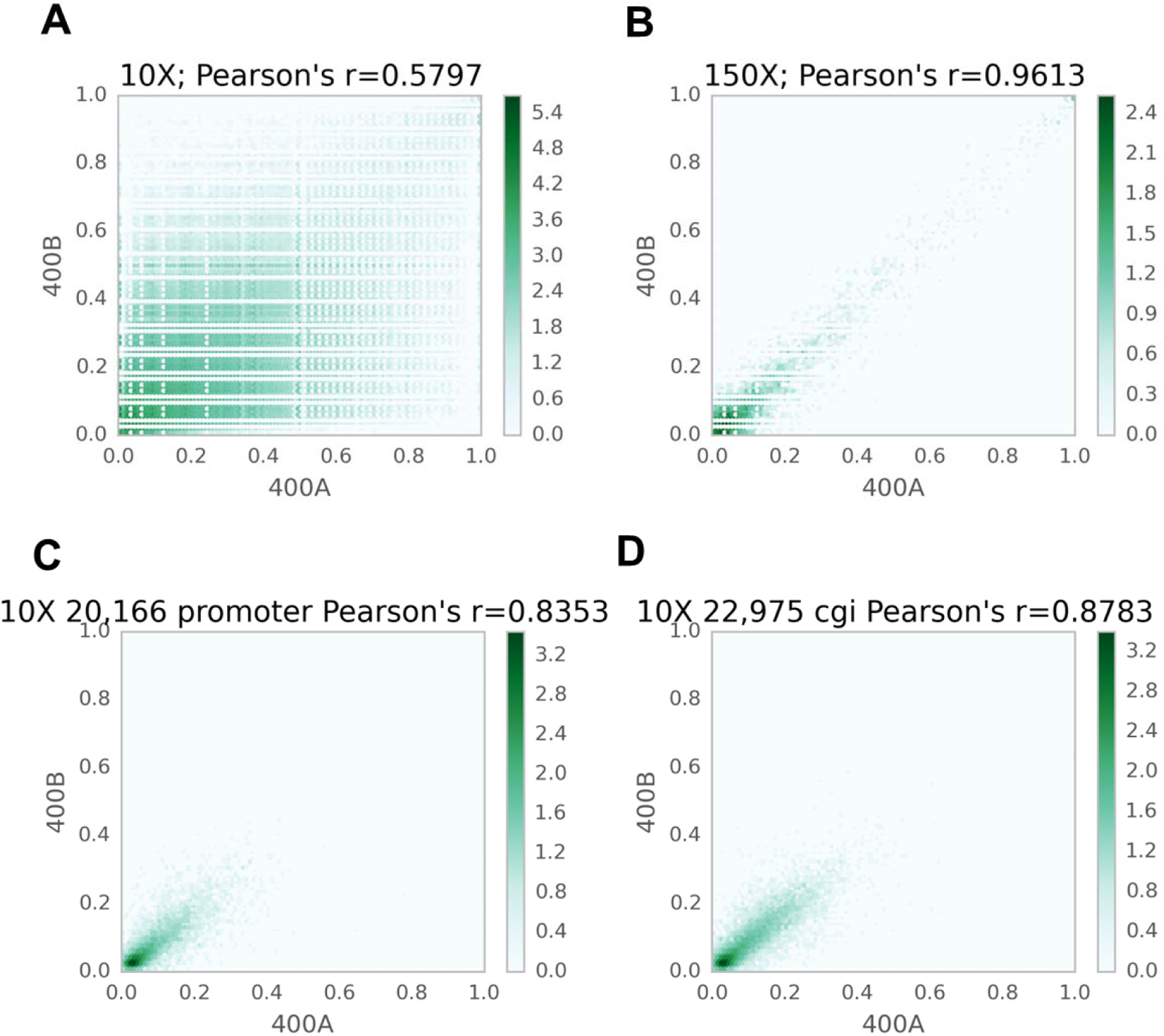
Genome-wide hydroxymethylation levels of the human brain. **(a)** The distribution of 5hmC and 5mC levels. **(b)** 5hmC and 5mC levels of various annotated regions. All CpG sites in both replicates have at least 10X coverage. Promoter was defined as 1kb upstream and downstream of TSS. TSS, exon, intron, CpG islands, LINE, SINE, LTR, and satellite were obtained from the UCSC genome browser track. Active Enhancer and Poised Enhancer were characterized previously^13^. 5hmC ratio was calculated using Mirror-RRBS data and mC ratio was calculated from RRBS data.

**Figure 4.**
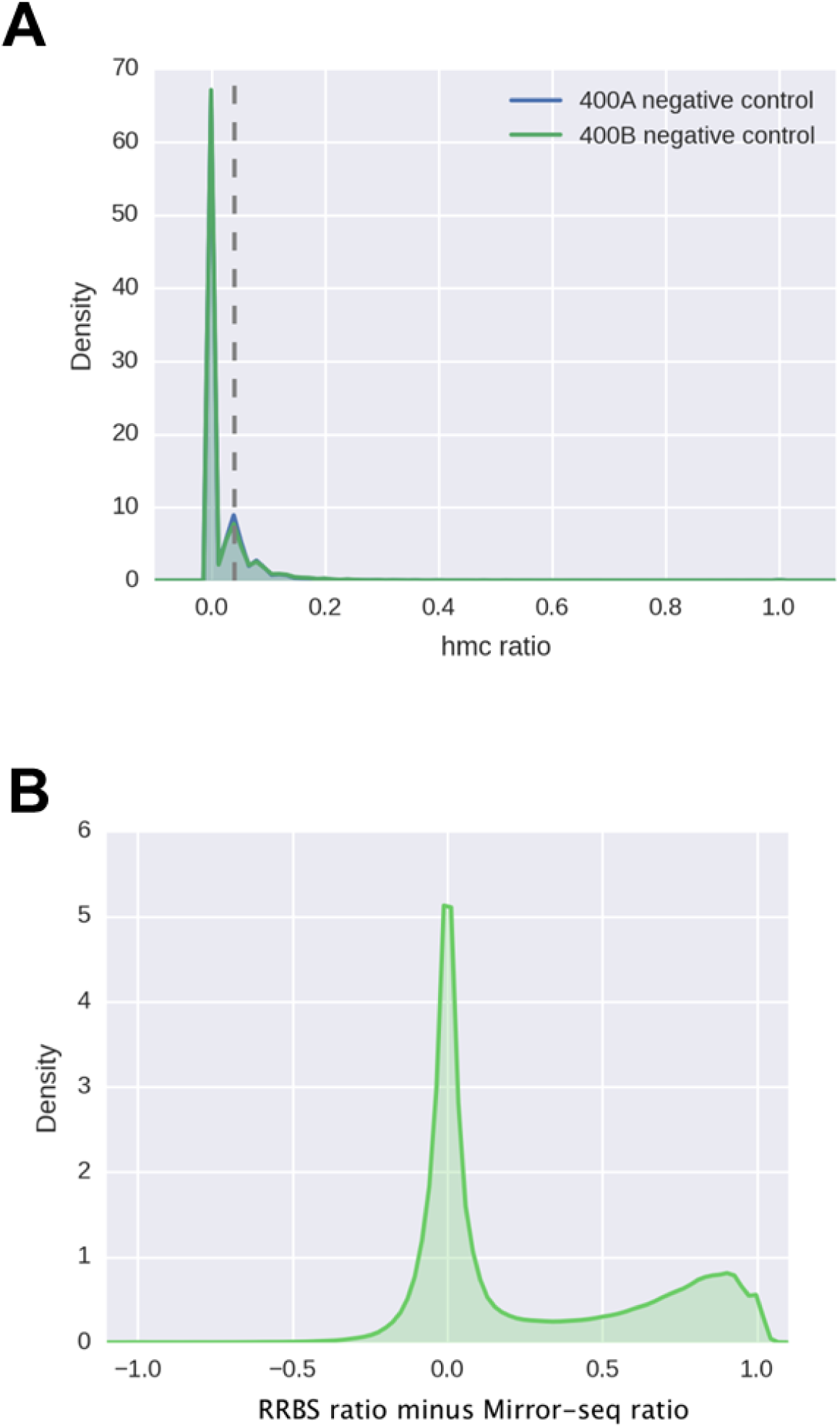
Evaluating the background level of Mirror-RRBS. **(a)** The distribution of false 5hmC levels detected by 400A and 400B negative control libraries, and the dotted line at 0.05 represented the 5hmC ratio where most CpG sites were falsely called. All CpG sites analyzed had at least 20X read coverage for both libraries. **(b)** The distribution of CpGs versus the difference between RRBS 5mC + 5hmC and Mirror-seq 5hmC levels. All CpG sites compared had at least 10X coverage in both libraries.

**Figure 5.**
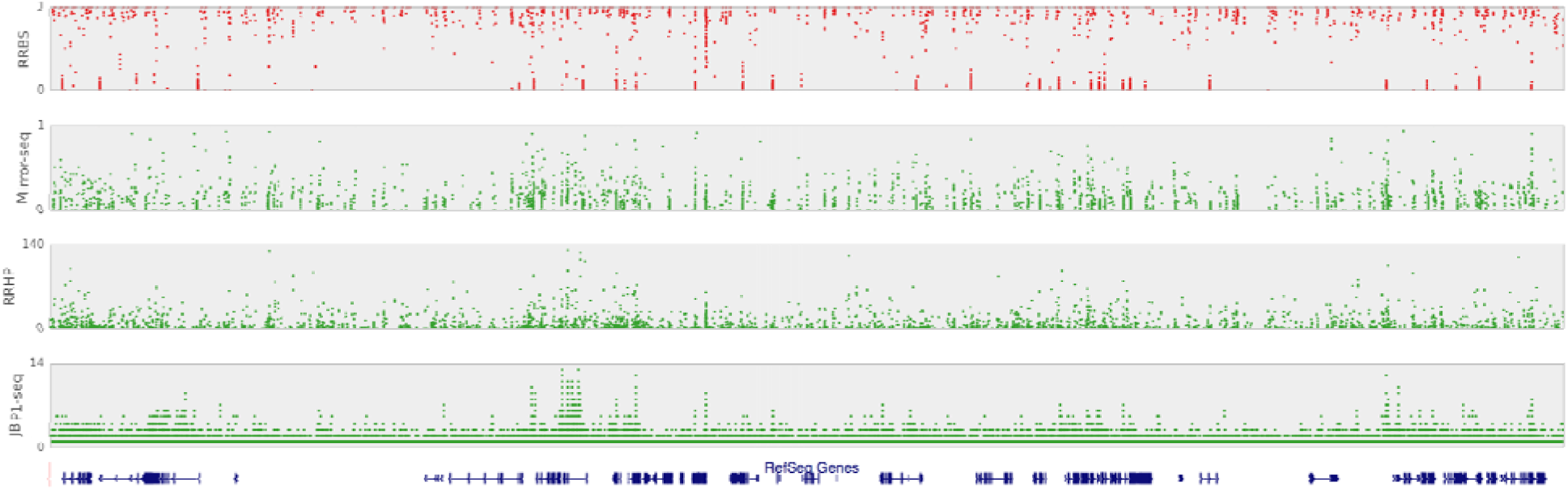
Comparison of 5hmC detection at hg19 chr4: 95,280,000-107,500,000 using various methods. RRBS detected both 5mC + 5hmC modifications, and Mirror-RRBS detected only 5hmC. RRHP is a positive-display method that determines the level of 5hmC by the number of tagged reads. JBP1-seq showed the relative enrichment of regions to background levels; higher 5hmC regions had higher folds of enrichment.

We also prepared two negative control libraries (400A negative and 400B negative) that follow the same library preparation but the βGT enzyme was omitted from the glucosylation reaction. Ideally, all CpG sites should be fully methylated by M.SssI and called as an unhydroxymethylated site. As expected, we found the majority of CpG sites had 5hmC levels under 5%, but we found a small portion of CpGs called as 100% hydroxymethylated (Fig. 4a). Further analysis and comparison to the common SNPs of dbSNP build 138 available from NCBI showed that those sites overlapped with SNPs in which CpG was altered into CpA. Other sites with high background were enriched in the DNA repeat region; we do not know the exact reason for the higher background levels in these regions, but it might be related to either structural variation or sequence specific features that inhibited M.SssI methylation. We sought to examine those sites and their neighboring regions for GC content, CpG density, and conserved sequence motifs, but similar to a previous study (Wu et al. 2014), we were unable to find any convincing associated trends. Interestingly, those sites are repeatedly observed in negative control samples from other tissues such as liver, kidney and spleen (Supplementary table 4), indicating there are hotspots for false 5hmC calling in Mirror-Seq and should be carefully examined and removed from analysis (Supplementary table 5).

Since traditional RRBS detects both 5mC and 5hmC, all of the methylation values called from traditional RRBS should be greater or equal to that of Mirror-RRBS. Therefore, to evaluate the reliability of Mirror-RRBS, we subtracted the 5hmC level detected by Mirror-RRBS from the 5hmC + 5mC levels detected by traditional RRBS, and we found the majority of the CpG sites met this criterion (Fig.4b). For sites with higher value in Mirror-Seq than in RRBS, we found only 4% of them are statistically different, which meant most of the sites in this category were not reliably called from either traditional RRBS, Mirror-RRBS, or both under the current sequencing depth (Supplementary fig. 6). If we excluded those sites, most of the CpG sites had both 5hmC and 5mC coexisting, but overall, 5hmCs occur at a relatively lower frequency with a median of 6% in comparison to 24% methylation (Figure 2).

**Figure 6.**
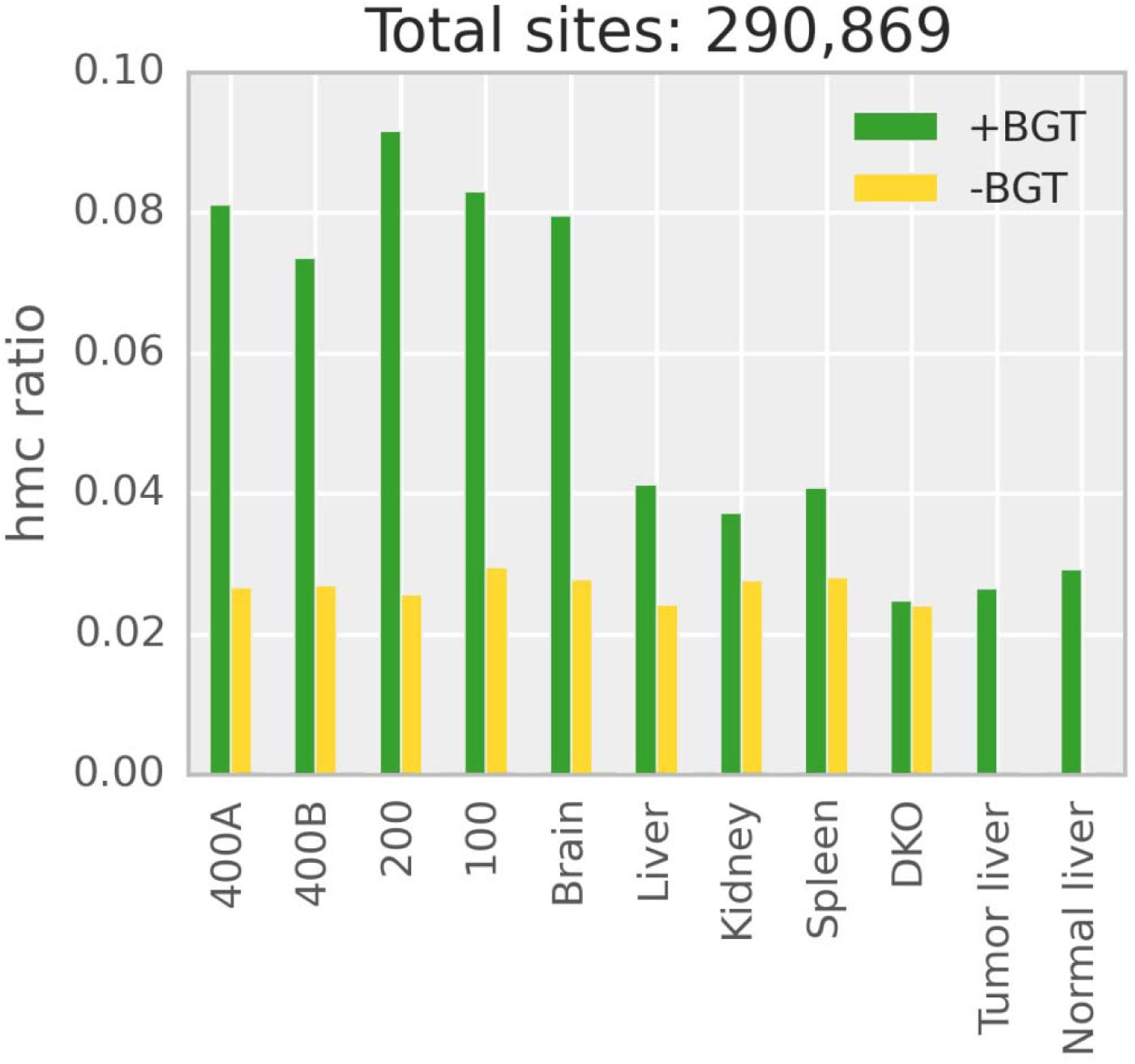
The global 5hmC level of the overlapping CpGs (> 10X read coverage) between different tissues.

To validate our method, we compared Mirror-RRBS data with the RRHP data which we had published previously (Petterson et al. 2014). RRHP is a positive display method that counts the tag of the sequencing reads to infer the 5hmC abundance. The principal behind the methodology is similar to glucMS-qPCR, but it is adapted for genome-wide analysis. We checked all of the overlapped sites between RRHP and Mirror-RRBS and found a correlation coefficient above 0.5 using Spearman analysis (Supplementary fig. 7a), which was similar to the correlation observed between oxBS-seq and glucMS- qPCR (Bhattacharyya et al. 2013). Since 5hmC is frequently found to be asymmetrical at the CpG site, enzymatic based methods such as RRHP and glucMS-qPCR may provide limited information since it is not clear whether 5hmCs are located exclusively on one of the two strands or present on both simultaneously, which could generate differential digestion patterns. That may explain the relatively low concordance between enzymatic- and bisulfite sequencing-based methods, but the general trend was consistent. In addition, we also compared the Mirror-RRBS data with previously published JBP1-Seq data (Cui et al. 2014); again, we also observed a similar correlation (Supplementary fig. 7b). Using all three methods for 5hmC detection, a locus-specific comparison was done for a 12 Mb genomic region in chromosome 4 (Fig. 5) that had been previously analyzed (Wen et al. 2014). RRBS data showed that 5mC was relatively constant in this region, but 5hmCs, on the other hand, were abundant in only a few enhancer regions, as confirmed by all three methodologies.

Finally, using Mirror-RRBS, we examined 5hmC abundance in several human tissues, including liver, kidney and spleen, as well as a double DNA methyltransferase knockout (DKO) cell line and paired liver tumor samples. Based on the overlapping sites detected for the different tissues, we found that the brain had the highest average 5hmC level, followed by liver, spleen and kidney (Fig. 6). This was consistent with the general trend observed previously (Li et al. 2011). In contrast, the DKO cell line showed a baseline level of 5hmC that was indistinguishable from the negative control sample. Since the DKO cell line had a double knockout of DNA methyltransferases (DNMTs), the low level of 5hmC was expected since the 5mC precursor was missing. In agreement with previous studies, we also found 5hmC patterns that were tissue specific, and some of the genomic loci with unique 5hmC patterns are shown in Supplementary figure 8 and a list of top tissue specific sites are in Supplementary table 6. Interestingly, we found that the tumor liver and its adjacent normal liver samples behaved quite similarly to the DKO cell line, and both of them had much lower 5hmCs than the healthy liver tissue.

Unsupervised clustering using all the 5hmC sites also grouped the paired tumor samples with the DKO cell line. It was known 5hmC underwent global depletion in tumor samples and such decrease was accompanied with the tumor progression (Yang et al. 2013). The low levels of 5hmC may be reflected by the higher grade of the tumor sample (grade IIIA), and while we expect the adjacent normal liver to have 5hmC levels similar to the healthy liver sample, it is possible that the adjacent tissues may have experienced 5hmC depletions earlier than its pathological changes. Pairwise comparison between liver tumor and adjacent normal tissues identified 2,107 significantly different 5hmC sites (Supplementary table 7). Most of them were not overlapping with the methylation changes, indicating 5hmCs were regulated independently from 5mC.

## DISCUSSION

In summary, we have developed an alternative, novel method, Mirror-Seq, for quantitative detection of 5hmC at single-base resolution. Compared with TAB-seq or oxBS-seq, Mirror-Seq does not involve oxidation steps, and the enzymes used in the preparation are efficient and easily accessible. Since it does not utilize oxidation steps that can result in the degradation and loss of DNA, this method requires much less genomic DNA input, allowing the accommodation of clinical and precious samples. This method is also suitable for whole genome hydroxymethylation analysis as well, but sequencing depth should be taken into consideration as discussed previously. Admittedly, Mirror-Seq is limited to the detection of 5hmC in the CpG context. Although non-CpG methylation is present in human, it has been found that over 99% of 5hmCs exist in the CpG context (Yu et al. 2012). Using human brain DNA as a testing material, we have extensively evaluated this method in terms of the reproducibility, sensitivity, specificity and reliability. We also examined 5hmC abundance in different human tissues such as liver, spleen, kidney as well as a paired tumor samples. The results were consistent with the previous findings and further validate the 5hmC detection capacity of Mirror-Seq.

## METHODS

### 100-mer dsDNA control

The following 100-mer oligonucleotide was synthesized with a single hydroxymethylated-CpG site: 5’-GTTACGCAAAGGAAGACCAAGAGTATGGTGGTCACTCGACCC AAAGGAATAT/i5HydMe-dC/GGTCCGGGTACGACAGGAGTCAGATAGAACCAGACCCAGAGGAGGCG-3’ (IDT).

The reverse complementary of the 100-mer oligo was synthesized by a primer (5’- CGCCTCCTCTGGGTCTGGTTCTATCT-3’) with ZymoTaq 2X PreMix (Zymo Research) using the thermal profile 95°C for 10 min, 50°C for 15 min, 72°C for 45 min and hold at 4°C. The dsDNA oligo was purified using the DNA Clean & Concentrator-5 (Zymo Research) and 500 ng of the dsDNA oligo was glucosylated with 6 U 5hmC glucosyltransferase (Zymo Research) and 100 μM uridine diphosphoglucose (UDPG) at 37°C for 12 hrs. The reaction was cleaned using the Oligo Clean & Concentrator (Zymo Research), and the purified oligo was CpG methylated using 4 U M.SssI methylase and 600 μM SAM (Zymo Research) at 30°C for 4 hrs and heat-inactivated at 65°C for 15 mins. After the incubation, the oligo was bisulfite converted by adding 130 μl of the conversion reagent from the EZ DNA Methylation – Lightning kit (Zymo Research) directly to the reaction and processed according to the manufacturer’s protocol. The newly synthesized strand was then selectively amplified with ZymoTaq 2X Premix and 10 μM of strand specific primer: Fwd: 5’ CAAAAAAAGACCAAAAATATAATAATCACTC and Rev: 5’ GTTTTTTTTGGGTTTGGTTTTATTTGATTTTTG. The reaction was first incubated at 95°C for 10 mins followed by 30 cycles of 95°C for 30 sec, 50°C for 30 sec, and 72°C for 30 sec and a final extension at 72°C for 1 min. The amplified DNA was purified using DNA Clean & Concentrator-5 (DCC-5), and 400 ng of DNA was digested with 20 U BspDI for 1 hr at 37°C and then visualized on a 4% agarose gel/TAE with ethidium bromide.

### Spike-in controls

The spike-in controls were generated by amplifying short regions (approximately 175 – 191bp) from *Escherichia coli K12* with two CCGG sites approximately 155bp apart (Supplementary Table 1). The spike-ins had a single 5hmCpG site, and it is located at the internal cytosine of the upstream CCGG site. This hydroxymethylated site was generated by synthesizing the forward primer to contain the 5hmC site at the desired position. The spike-in controls were amplified using ZymoTaq 2X Premix using either the forward primer with or without the 5hmC modification at the CCGG site. The thermal cycler condition was 95°C for 10 mins, 30 cycles at 95°C for 30 sec, 58°C for 30 sec, and 72°C for 1 min. The amplicons were purified using DCC-5 and quantified by quantitative PCR. The 5hmC and C amplicon of the same sequence were mixed to the desired 5hmC%, which was then used as a spike-in.

### DNA samples

Human brain, liver, kidney, and spleen genomic DNA as well as DKO cell line genomic DNA was obtained from Zymo Research. The paired liver samples, a tumor and a normal adjacent tissue, were obtained from ILSbio, and genomic DNA was extracted using the ZR Genomic DNA Tissue Miniprep (Zymo Research).

### RRBS library preparation

Libraries were generated by digesting and adapterizing 400ng of human brain genomic DNA (Zymo Research) with 40 U MspI (NEB), 400 U T4 DNA Ligase (NEB), 100 μM ATP, and 200 nM of modified Illumina P5 and P7 TruSeq adapters at 21°C for 6 hrs. After incubation, adapters were extended with 2.5 U GoTaq DNA polymerase (Promega) and 300 μM 5-methylcytosine dNTP mix (Zymo Research) at 74°C for 30 mins, and then the libraries were purified using DCC-5. The libraries were size-selected on a 2.5% NuSieve/Agarose gel with ethidium bromide into small (150-250 bp) and large (250-350 bp) fragments and purified using the Zymoclean Gel DNA Recovery Kit (Zymo Research). The libraries were then bisulfite converted using the EZ DNA Methylation – Lightning Kit (Zymo Research) and amplified with 200 nM P5 and P7 barcoding primers in OneTaq Hot Start 2X Master Mix (NEB) with an initial denaturation of 94°C for 30 sec, 14 cycles of 94°C for 30 sec, 58°C for 30 sec, and 68°C for 1 min, and a final extension at 68°C for 5 mins. The amplified libraries were purified with the DCC-5 kit, and equal molarities of the small and large fragments were then pooled for each sample. Six libraries were multiplexed using equal masses and sequenced on Illumina HiSeq 2000 using 50 bp paired-end reads (Illumina).

### Mirror-RRBS library preparation

The initial digestion and adapterization of genomic DNA was prepared using the described procedure above. After adapter extension and purification, a single strand was then synthesized using OneTaq Hot Start 2X Master Mix (NEB) to mirror the parental strand by utilizing a primer complementary to the P7 adapter that has methylated cytosines: 5’- GTGA/iMe-dC/TGGAGTT/iMe-dC/AGA/iMe-dC/GTGTG/iMe-dC/T/iMe-dC/TT/iMe-dC//iMe-dC/GAT/iMe-dC/*T – 3’ (IDT). The single-strand extension reaction was incubated at 95°C for 3 mins, 55°C for 2 mins, and 68°C for 16 mins and then purified using the DCC-5 kit. 5hmC sites were then glucosylated with 8 U 5hmC glucosyltransferase (Zymo Research) and 100 μM UDPG at 37°C for 2 hrs and heat-inactivated at 65°C for 15 mins, and the DNA was purified using DCC. The libraries were methylated with 8 U CpG Methylase (Zymo Research) and 600 μM SAM at 30°C for 4 hrs and heat-inactivated at 65°C for 15 mins. The libraries were size-selected as described above followed by bisulfite conversion using the EZ DNA Methylation – Lightning Kit (Zymo Research). Mirror-RRBS libraries were amplified with 200 nM P5 and P7 barcoding primers in OneTaq Hot Start 2X Master Mix (NEB) with an initial denaturation of 94°C for 30 sec, 15 cycles of 94°C for 30 sec, 58°C for 30 sec, and 68°C for 1 min, and a final extension at 68°C for 5 mins. The amplified libraries were then purified, pooled, and sequenced using the same parameters as described above.

### Bioinformatics analysis

For Mirror-RRBS analysis, Illumina’s TruSeq adapter sequence and low quality bases at the end of reads were trimmed off from the Fastq files by Trim Galore! v0.2.3. A Python script trimmed off filled-in nucleotides after Trim Galore!, and Bismark v0.11.1 with — non_directional argument was used to align reads to hg19 human reference genome. A Python script was used to perform hydroxymethylation calling by first calculating the hydroxymethyation ratio (*r*) as T/(C+T) at each called CpG site and then assigning 1-*r* to the cytosine site of the other strand. For RRBS samples, Trim Galore! and Bismark were also used to do trimming and alignment under the same parameters as Mirror-RRBS samples. A Python script was used to remove filled-in nucleotides, and methylation ratio was calculated as C/(C+T) at each called cytosine.

### Tissue-specific hydroxymethylated sites

To find tissue specific 5hmC sites of tissue *s* compared with other tissues *S*, one sample t-test was performed using hydroxymethylation ratios of *s* and *S*. Sites with p-value smaller than 0.05 and the absolute difference between 5hmC level of *s* and the average 5hmC level of *S* greater than 0.3 were filtered. After sorting by absolute hydroxymethylation difference, top 1000 sites were selected as tissue specific sites (Supplementary table 6). For comparison of the liver tumor and its adjacent normal tissue, overlapping sites with more than 0.3 absolute differences in hydroxymethylation ratio were selected. P-values were generated by Fisher’s exact using hydroxymethylated read number and non-hydroxymethylated read number of both samples. Sites with a p-value larger than 0.05 were filtered out.

## DATA ACCESS

The sequencing data from this study have been submitted to NCBI’s Gene Express Omnibus (GEO) (http://www.ncbi.nlm.nih.gov/geo) under accession no. GSE69272. The following link has been created to allow review of record GSE69272 while it remains in private status (http://www.ncbi.nlm.nih.gov/geo/query/acc.cgi?token=gnatcamkhvyznaf&acc=GSE69272).

## ACKNOWLEDGMENTS

We thank P. Shi, A. Kim, and K. Booher for assistance in data acquisition and analysis, and M. Van Eden, J. Claypool, and L. Cui for helpful discussion.

## AUTHOR CONTRIBUTIONS

X.S. and X.J. conceived the idea and X.S. designed the study. D.T. constructed libraries and performed sequencing experiments. T.H.C performed bioinformatics processing of sequencing data and all subsequent statistical analyses. X.S. and D.T. wrote the manuscript, and all authors have read and approved the final manuscript.

